# Combined Developmental Toxicity Of Cyhalofop-butyl And Quizalofop-p-ethyl On The Zebrafish (Danio rerio) Embryos

**DOI:** 10.1101/2022.10.04.510810

**Authors:** Li Liu, Dongmei Wang, Ping Li, Huan Zhao

## Abstract

Aryloxyphenoxypropionate herbicides have the characteristics of high efficiency, low toxicity, and safety to subsequent crops, and occupy an important position in the world herbicide market. Cyhalofop-butyl and quizalofop-p-ethyl are two representative herbicides, which are widely used in weed control. However, there is limited information on their combined toxicity to aquatic organisms. In this study, the developmental toxicity of cyhalofop-butyl and quizalofop-p-ethyl exposure in combination on zebrafish embryos was valuated to better understand the interaction between the that. The 96 h-LC_50_ (50% lethal concentration) of cyhalofop-butyl and quizalofop-p-ethyl on zebrafish embryos were 0.637 mg·L^−1^ and 0.248 mg·L^−1^, respectively. The combined effect of cyhalofop-butyl and quizalofop-p-ethyl was an antagonistic effect, and the 96 h-LC_50_ of zebrafish embryos was 1.043 mg·L^−1^. Morphologically distinct pericardial edema and yolk cysts were observed after combined exposure, with significant effects on body length and heart rate in zebrafish embryos. At the same time, the mRNA levels of gene related to apoptosis and cardiac development also changed significantly. Therefore, we speculate that changes in genes related to apoptosis and cardiac development should be responsible for the abnormal development during embryonic development following co-exposure of cyhalofop-butyl and quizalofop-p-ethyl.

**Highlights:**

- Combined exposure caused deformities in zebrafish.
- Combined exposure caused apoptosis in zebrafish.
- Combined exposure altered the expression of apoptosis and cardiac-related genes in zebrafish.

## 1. Introduction

Like most other water pollutants, pesticides can enter hydrological systems through diffusion pathways (non-point sources) from specific release points (point sources) and deposition and surface runoff (Chang et al., 2020; Hladik et al., 2014). While aquatic organisms are often exposed to a variety of pollutants, a variety of pesticides combine to produce different toxic effects. However, most of the current studies have only assessed the effects of a single compound on aquatic organisms (Zhu et al., 2015). Therefore, when assessing the harm of pesticides to aquatic organisms, the effect of combined exposure to pesticides should be fully considered. Aryloxyphenoxypropionate compounds are a class of highly active herbicides, and their mechanism of action is to inhibit the activity of acetyl-CoA carboxylase (ACCase) in gramineous plants, and occupy an important position in the world herbicide market (Cummins and Edwards, 2004; Kukorelli et al., 2013). Studies have shown that the combined toxicity of fenoxaprop-ethy and cyhalofop-butyl was evaluated, and it was found that these two herbicides could reduce the content of chlorophyll a and b in duckweed (Fan et al., 2006). The fenoxaprop-ethy toxicity is highly toxic and has certain genotoxicity to grass carp (Chen et al., 2006). Therefore, it is of great significance to strengthen the research on the toxicology of aryloxyphenoxypropionate herbicides.

Cyhalofop-butyl is a new type of highly effective aryloxyphenoxypropionic acid herbicide widely used for weed control in paddy fields (Zuo et al., 2016). Unfortunately, cyhalofop-butyl in rice fields will inevitably cause this herbicide to enter the aquatic environment and disturb the balance of farmland ecosystem (Xia et al., 2018; Yuan et al., 2019). Previous studies have shown that 7-day exposure of 0.662 mg/L cyhalofop-butyl to *misgurnus anguillicaudatus* will lead to vacuolation and swelling of liver cells, and with the extension of exposure time, cell lysis, nuclear deformation and nuclear pyknosis will also appear (Shang et al., 2019).

Quizalofop-p-ethyl is a selective transport agent for stems and leaves of aryloxyphenoxypropionate (Li et al., 2012). Some studies have pointed out that quizalofop-p-ethyl will pollute water bodies, possibly affecting the aquatic ecological environment and aquatic organisms (Belfroid et al., 1998). At present, the biotoxicity of quizalofop-p-ethyl, such as genotoxicity (Mustafa and Arikan, 2008), reproductive toxicity (Zhu et al., 2017) and hepatotoxicity (Elefsiniotis et al., 2007) has been reported. For example, some studies have found that quizalofop-p-p-ethyl can affect the biological indicators such as molting and reproduction of the parent daphnia magna (Yang-Guang et al., 2013). The research team also found that the acute toxicity of quizalofop-p-p-ethyl to zebrafish adults was high, and both of them affected the activity of ATPase (Yang-Guang, 2012). Zhu et al. (2017) revealed for the first time that quizalofop-p-ethyl as an endocrine disrupting chemical (EDC) destroyed the endocrine system of zebrafish. In addition, quizalofop-p-ethyl aggravated cardiotoxicity due to inflammatory reaction, which affected the development of zebrafish embryos (Zhu et al., 2022).

Because zebrafish has the advantages of small size, low feeding cost, strong reproduction ability, transparent embryo, short test cycle and high similarity between genes and human genes, it is widely used in water quality testing experiments for environmental pollutant toxicity assessment (Howe et al., 2013). The zebrafish model used to study human diseases is widely accepted in various fields, including developmental genetics, toxicology, cancer and regeneration (Shaukat et al., 2011; Wolfram and Sadler, 2018; Yan et al., 2016). In this study, the acute lethal and developmental toxicity effects of cyhalofop-butyl, quizalofop-p-p-ethyl and binary mixtures on zebrafish embryos were detected. The purpose of this study is to reflect the potential threat of joint toxicity of pesticides to aquatic organisms, so as to provide necessary basic data for comprehensive assessment of pesticide compound pollution risk.

## 2. Materials and methods

This study was approved by the research ethics from Chongqing University. The study was carried out under the Guidance of the Care and Use of Laboratory Animals in China.

### 2.1 Materials and reagents

98% cyhalofop-butyl (CAS: 122008-85-9) and 95% quizalofop-p-ethyl (CAS: 100646–51–3) was obtained from Anhui Futian Agrochemical CO., Ltd and Shandong China Jingbo Agrochemicals CO., LTD. Preparation of drug exposure stock solution with acetone AR, and all reagents used in the study are analytical grade.

### 2.2 Maintenance of zebrafish

The parental zebrafish were AB wild type. In the zebrafish culture system, the temperature was 27±1◻, and the light-dark period was 14 h:10 h. During the breeding period, the mortality rate was kept below 5%, and the shrimp were fed twice a day. Embryo acute toxicity and developmental toxicity tests were performed according to the previous reported method (Mu et al., 2016).

### 2.3 Toxicity test of zebrafish

#### 2.3.1 Acute toxicity test of cyhalofop-butyl and quizalofop-p-ethyl

In the acute toxicity experiment, embryos within 2 hours after fertilization (hpf) were collected according to the OECD standards (OECD, 1992) and exposed to cyhalofop-butyl and quizalofop-p-ethyl 96 hpf. The combined exposure was based on the LC_50_ values of single cyhalofop-butyl and quizalofop-p-ethyl determined, and a ratio of toxicity units of 1:1 was used in the experimental design. All test solutions were made in 150ml small beakers, each containing 100 ml of exposure solution for 20 larvae. There were three replicates for each concentration group. The recombinant water without cyhalofop-butyl and quizalofop-p-ethyl was used as a blank control (0 mg/L), and a solution with the same acetone content as the highest concentration was arranged as the solvent control, see Table 1 for the specific concentration setting. The deformity and death of larvae were observed every 24 h, and the dead individuals were removed in time. The LC_50_ was calculated by replacing 3/4 volume of liquid every 24 h. Combined toxicity assessment was performed according to Marking’s Additional Index (AI) method for combined effects of aquatic toxicology (Marking, 1984). The formula is as follows:

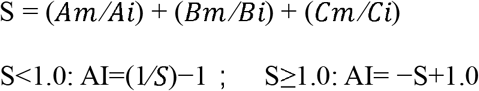

S: total biological activity. A, B and C: the compounds in the mixture, i: individual LC50 value for A, B, or C. m: LC_50_ of A, B, or C in mixture. The AI value can be obtained by substituting the calculated S value into the formula. When AI < 0, the joint toxic effect of the mixture is antagonistic, when AI=0, it is additive, and when AI > 0, it is synergistic, and the larger AI value, the stronger the synergistic effect.

**Table 1.** Concentration of cyhalofop-butyl and quizalofop-p-ethyl in acute toxicity

| Number | Cyhalofop-butyl/(mg/L) | Quizalofop-p-ethyl/(mg/L) | Cyhalofop-butyl+Quizalofop-p-ethyl/(mg/L) |
| --- | --- | --- | --- |
| 1 | 1.52 | 0.94 | 1.096+0.427 |
| 2 | 1.01 | 0.52 | 0.911+0.355 |
| 3 | 0.68 | 0.29 | 0.758+0.295 |
| 4 | 0.45 | 0.16 | 0.637+0.248 |
| 5 | 0.30 | 0.09 | 0.529+0.206 |
| 6 | 0.20 | 0.05 | 0.440+0.171 |

### 2.4 Embryonic development toxicity of co-exposure

According to the LC_50_ of cyhalofop-butyl and quizalofop-p-ethyl, two concentrations and one solvent control were set, and each concentration was set in triplicate. The concentrations of cyhalofop-butyl were set to 0.166 and 0.333 mg/L, and the concentrations of quizalofop-p-ethyl were set to 0.065 and 0.129 mg/L. The combined exposure concentration of cyhalofop-butyl + quizalofop-p-ethyl was set to 0.166 mg/L + 0.065 mg/L, 0.333 mg/L+0.129 mg/L. For each concentration, 200 normally developing embryos were randomly selected and placed into a 1000 mL beaker containing 600mL of exposure solution. The experimental group and control group were exposed to 27±1 ◻ incubator for 96 h, the light-dark cycle was 14 h∶ 10 h, and the reagent was changed every 24 h. The spontaneous movement of 24 hpf, hatching rate of 96 hpf, heart rate, body length of larvae (Aigo ge-5, China) and deformity rate (Olympus BH-2 dissecting microscope) were recorded. At the end of exposure, 50 zebrafish larvae at each concentration were randomly selected for the determination of apoptosis enzyme activity and 30 zebrafish larvae for the determination of genes related to apoptosis and heart development.

### 2.5 Measurement of apoptosis enzyme activity

The caspase-3, caspase-9 activity and the total protein of samples was measured by caspase assay kit and BCA protein kit according to the manufacturer’s method (Beyotime Institute of Biotechnology, Haimen, China).

### 2.6 Gene expression analysis

RNA was extracted from each replicated zebrafish embryo using Trizol (Tiangen Biotech, China) and reverse transcribed by a quant cDNA (First strand complementary DNA) synthesis RTase Kit (Tiangen Biotech, China). Quantitative real-time polymerase chain reaction (qRT-PCR) was performed SuperReal PreMix Plus kit (Tiangen Biotech, China) by ABI 7500 Q-PCR system (Applied Biosystems, USA). Reaction system conditions and primer sequences are detailed in the Supplementary Material. The housekeeping gene beta-actin (β-actin) was used as the reference gene and was quantified by 2^−ΔΔ Ct^ method. All measurements were performed in triplicate.

### 2.7 Statistical analysis

SPSS 22.0 (SPSS, Chicago, IL, USA) was used to analyze the experimental data. One-way analysis of variance was used for significance analysis, and Dunnett test was used to evaluate the significant differences between different groups. P< 0.05 was considered statistically significant, and values were expressed as mean ± standard deviation (SD).

## 3. Results

### 3.1 Acute toxicity of cyhalofop-butyl and quizalofop-p-ethyl

The results showed that the toxicity of cyhalofop-butyl and quizalofop-p-ethyl on zebrafish were middle toxicity, and the LC_50_ value of 96 h were 0.637 mg/L and 0.248 mg/L, respectively. When the toxicity ratio of cyhalofop-butyl and quizalofop-p-ethyl was 1:1, the AI value of 96 h of combined exposure was −1.086, which had an antagonistic toxicity effect (Table 2).

**Table 2.** Acute toxicity test of cyhalofop-butyl, quizalofop-p-ethyl and co-exposure on zebrafish embryos.

| Chemical | Regression equation | 96h-LC<sub>50</sub> (mg /L)<br>(95% CL<sup>a</sup>) | R<sup>2b</sup> | AI | Effect<br>evaluation |
| --- | --- | --- | --- | --- | --- |
| ( Cyhalofop-butyl ) | y=0.634+3.243x | 0.637<br>( 0.551~0.728 ) | 0.937 | -1.086 | Antagonistic<br>effect |
| ( Quizalofop-p-ethyl ) | y=1.496+2.469x | 0.248<br>( 0.212~0.291 ) | 0.96 |  |  |
| Cyhalofop-butyl+Quizalofop-p-ethyl | y=-0.252+13.662x | 1.043<br>( 1.001~1.085 ) | 0.988 | — | — |

CL<sup>a</sup>: Confidence limit; R<sup>2b</sup>: Coefficient of determination

### 3.2 Embryonic development toxicity

Zebrafish embryos were exposed to cyhalofop-butyl, quizalofop-p-ethyl and their co-exposure for 24 h, the voluntary movement was inhibited (Figure 1-A). In the co-exposure treatment group, the inhibitory effect increased with the increase of exposure concentration. At 48 hpf, the single treatment group of cyhalofop-butyl and quizalofop-p-ethyl had no significant effect on the fetal heartbeat of zebrafish, while the co-exposure group significantly inhibited the fetal heartbeat of zebrafish. At 72 hpf and 96 hpf, the fetal heartbeat of zebrafish was significantly inhibited in all treatment groups except the low-concentration cyhalofop-butyl treatment group (0.166 mg/L) (Figure 1-B). After 96 hpF exposure, the hatchability of zebrafish treated with the highest concentrations of cyhalofop-butyl (0.333 mg/L) and quizalofop-p-ethyl (0.129 mg/L) in the co-exposure group was significantly inhibited (Figure 1-C). In addition, compared with the control group, the body length of zebrafish larvae in all treatment groups decreased with the increase of exposure concentration, and the body length of zebrafish larvae in the co-exposure group decreased significantly (Figure 1-D). Single and co-exposure of cyhalofop-butyl and quizalofop-p-ethyl induced pericardial edema and yolk sac edema deformity in zebrafish embryonic development (Figure 2 A-B). The results showed that the malformation rate in all treatment groups was significantly increased by dose effect. At 48 hpf and 96 hpf, the embryo malformation rate of zebrafish in the co-exposed high concentration treatment group significantly increased to 36.11% and 36.71%, respectively (Figure 2-C).

**Fig. 1.**
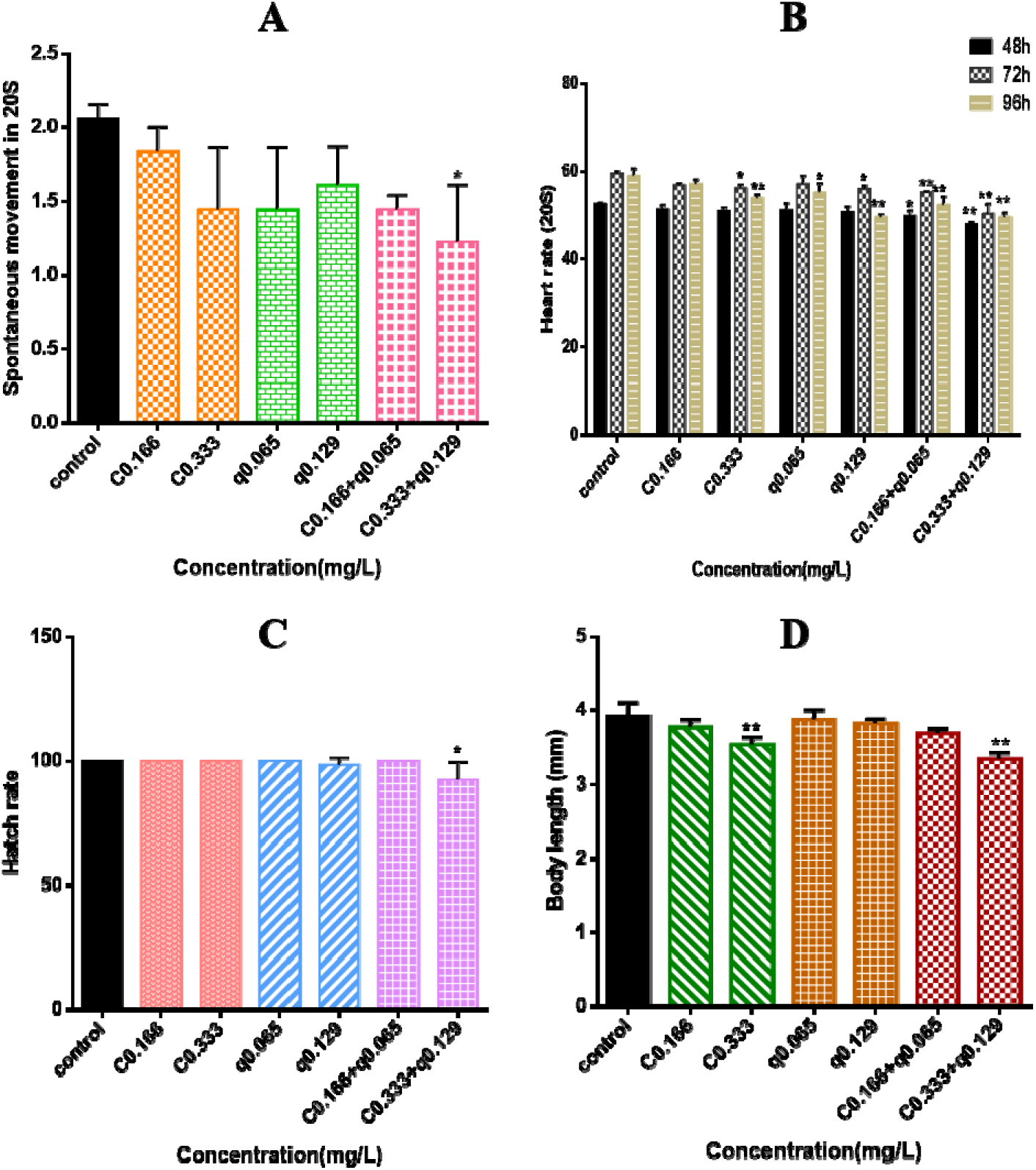
Developmental effects of cyhalofop-butyl, quizalofop-p-p-ethyl and binary mixtures on embryos. Asterisks denote significant difference between the treatments and control (determined by Dunnett post-hoc comparison, ^◻^*p* < 0.05; ^◻◻^*p* < 0.01). A: Spontaneous movement of zebrafish at 24hpf. B: Heart rate at each observation time C: Hatch rate at 96 hpf. D: Body length at 96 hpf C0.166: 0.166 mg/L cyhalofop-butyl; C0.333: 0.333 mg/L cyhalofop-butyl; q0.065: 0.065 mg/L quizalofop-p-p-ethyl; q0.129: 0.129 mg/L quizalofop-p-p-ethyl; C0.166+q0.065: co-exposure of cyhalofop-butyl + quizalofop-p-ethyl 0.166 mg/L + 0.065 mg/L; C0.333+q0.129: co-exposure of cyhalofop-butyl + quizalofop-p-ethyl 0.333 mg/L+0.129 mg/L).

**Fig. 2.**
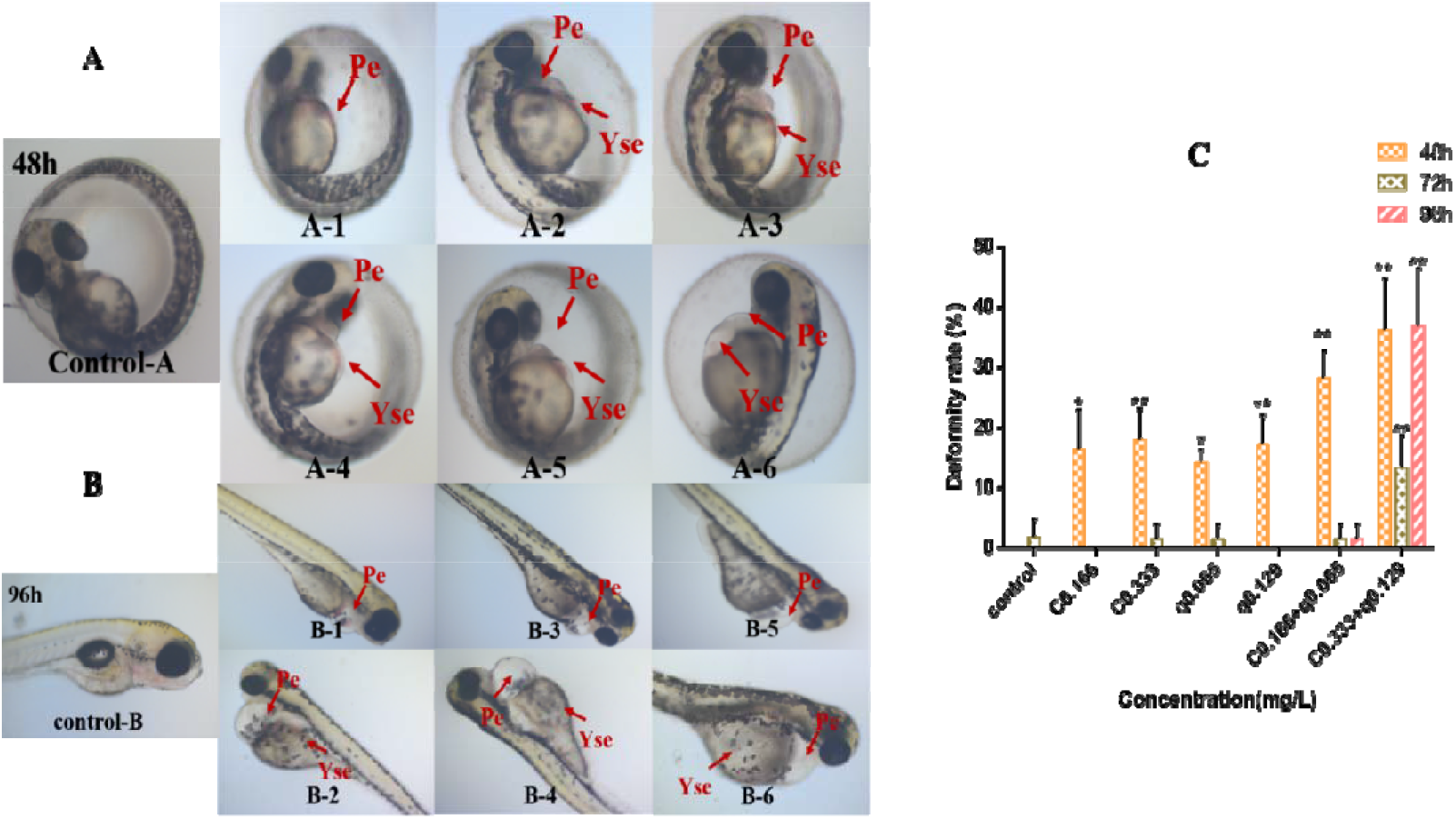
Malformations of embryos after exposure to cyhalofop-butyl, quizalofop-p-p-ethyl and binary mixtures. A: embryos with morphological deformation after exposure at 48 hpf. B: embryos with morphological deformation after exposure at 96 hpf (−1, 0.166 mg/L cyhalofop-butyl; −2, 0.333 mg/L cyhalofop-butyl; −3, 0.065 mg/L quizalofop-p-p-ethyl; −4, 0.129 mg/L quizalofop-p-p-ethyl; −5, co-exposure of cyhalofop-butyl + quizalofop-p-ethyl 0.166 mg/L + 0.065 mg/L; −6, co-exposure of cyhalofop-butyl + quizalofop-p-ethyl 0.333 mg/L+0.129 mg/L). C: Rate of deformity of embryos after exposure for 48,72 and 96 hpf. Arrows mark the different positions. Asterisks denote significant difference between the treatments and control (determined by Dunnett post-hoc comparison, ^◻^*p*< 0.05; ^◻◻^*p* < 0.01).

### 3.3 Apoptosis analysis

Compared with the control group, the activity of caspase-3 and caspase-9 in cyhalofop-butyl, quizalofop-p-ethyl and their co-exposure treatment groups increased with the increase of exposure concentration. The enzyme activities of caspase-3 and caspase-9 increased by 224.09% and 235.63% respectively in the treatment group with the highest concentration of co-exposure (Figure 3). There was no significant change in caspase activity of embryos in other treatment groups.

**Fig. 3.**
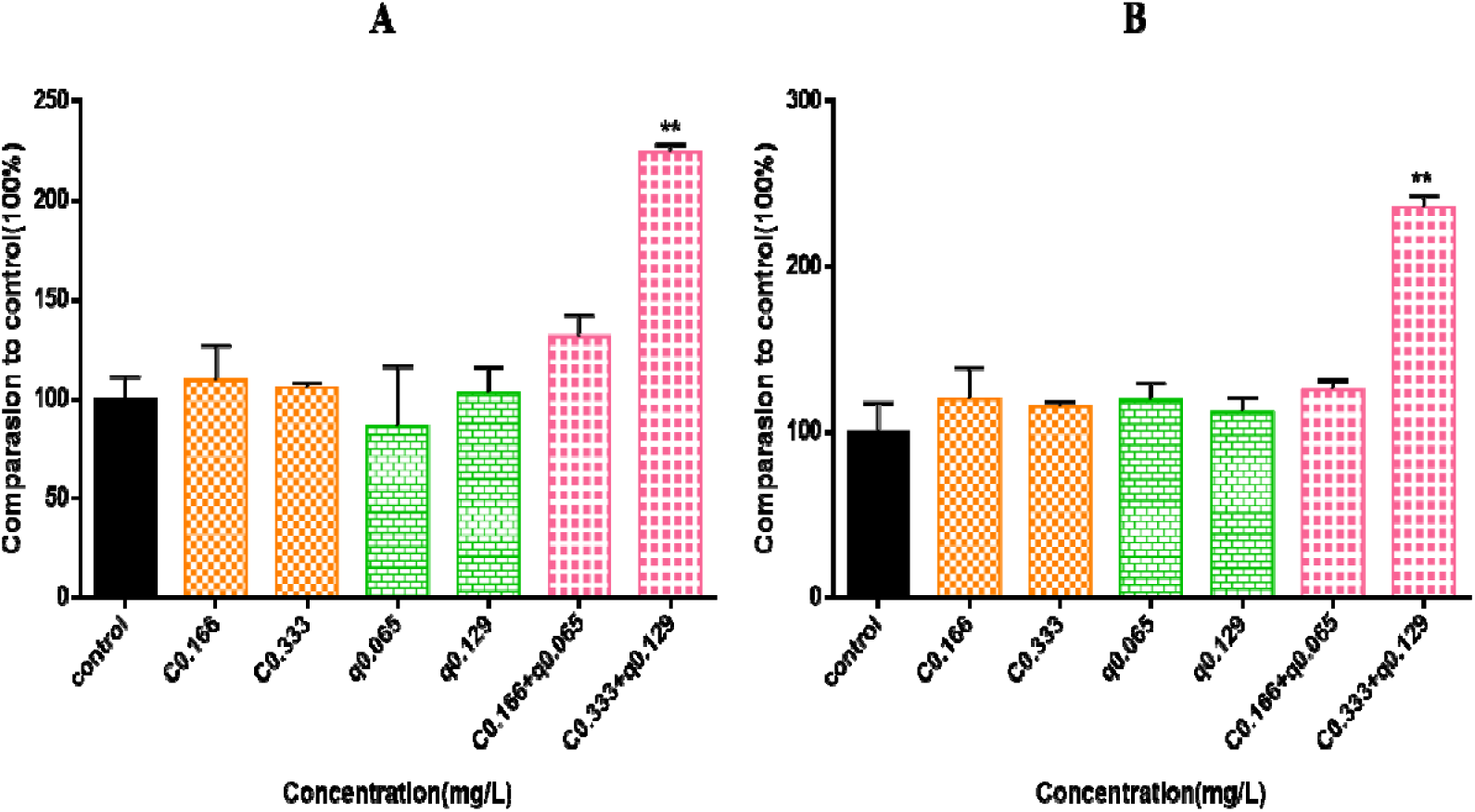
Cell apoptosis induced by cyhalofop-butyl, quizalofop-p-p-ethyl and binary mixtures. A and B: The activity of caspase-3 (A) and caspase-9 (B) of control and treated groups at 96 hpf. Asterisks denote significant difference between the treatments and control (determined by Dunnett post-hoc comparison, ^◻^*p* < 0.05; ^◻◻^*p* < 0.01).

At 96 hpf, the related genes apoptosis protein activating factor 1 (*apaf1*), Bcl-2 associated X protein (*bax*), B-cell lymphoma-2 (*bcl-2*), the relative expressions of cysteinyl aspartate specific protease 3 (*caspase-3*), cysteinyl aspartate specific protease 9 (*caspase-9*), murine double minute 2 (*mdm2*) and *p53* increased significantly in the co-exposure treatment group (Figure 4). Compared with the control group, *apaf1* (2.644 and 2.683 times), *mdm2* (2.968 and 6.510 times) and *p53* (3.349 and 2.36 times) were increased significantly with co-exposure (0.166+0.065 mg/L, 0.333+0.129 mg/L). *Bcl-2* was significantly expressed in quizalofop-p-ethyl (0.065 mg/L, 0.129 mg/L) and Co-exposure (0.166+0.065 mg/L, 0.333+0.129 mg/L) treatment groups, which were up-regulated by 1.892, 2.490, 2.119 and 3.279 times respectively.

**Fig. 4.**
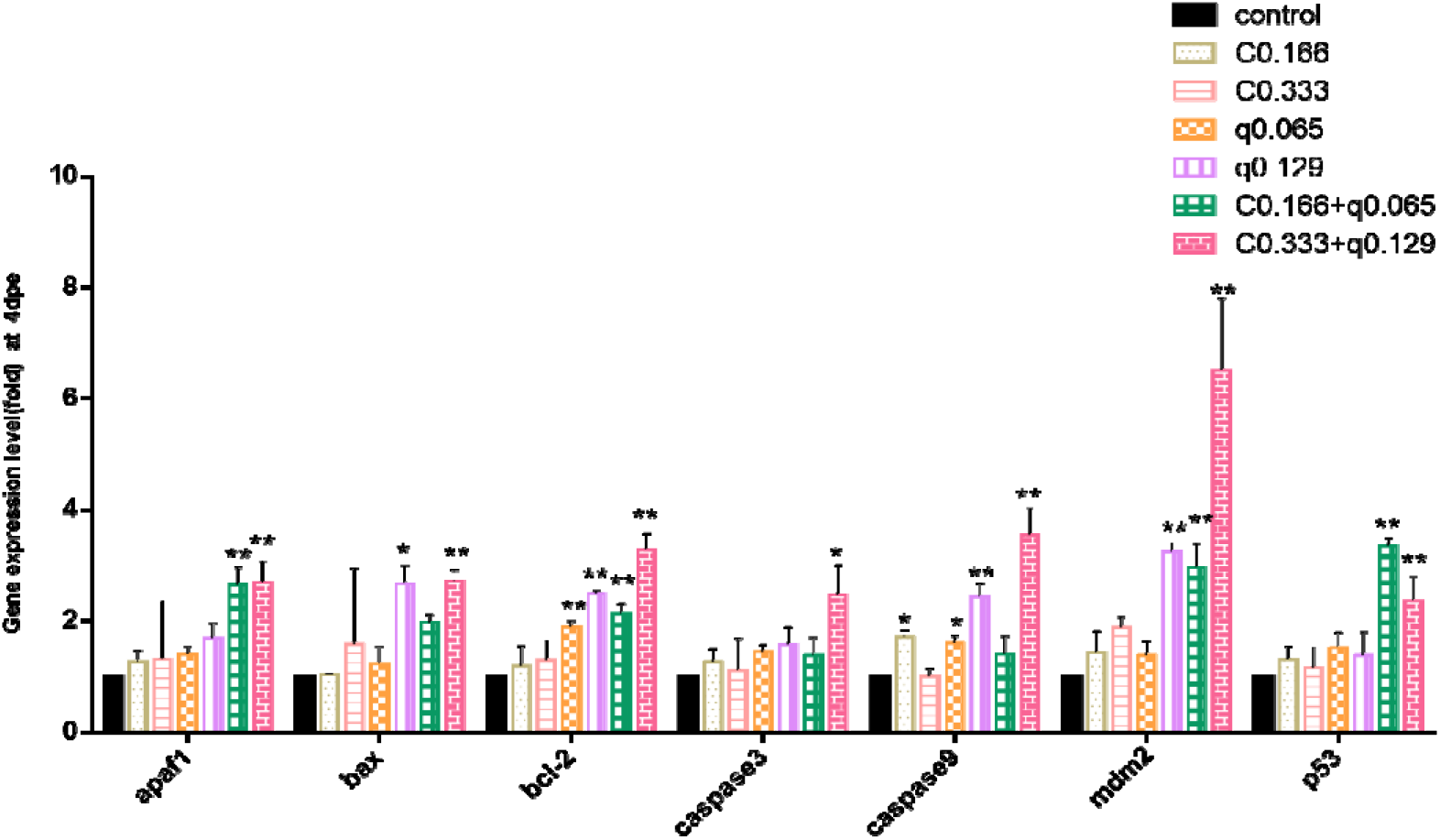
Transcription of apoptosis related genes in embryos after 96 h exposure to cyhalofop-butyl, quizalofop-p-p-ethyl and binary mixtures. Results are presented as mean ± SD of three replicate samples (determined by Dunnett post-hoc comparison, ^◻^*p* < 0.05; ^◻◻^*p* < 0.01).

### 3.4 The expression of genes related to heart development

At 96hpf, the expressions of *Nkx2.5*, *Tbx5* and *VEGF* were significantly up-regulated with the increase of concentration in all treatment groups, especially in the high co-exposure concentration group (0.333+0.129 mg/L). The expression of *Gata4* was down-regulated in cyhalofop-butyl and quizalofop-p-ethyl treatment groups, but it was significantly increased with the increase of concentration in co-exposure treatment group. The expression of *vmhc* gene was significantly induced in all treatment groups, and the up-regulation level increased with the increase of concentration (Figure 5).

**Fig. 5.**
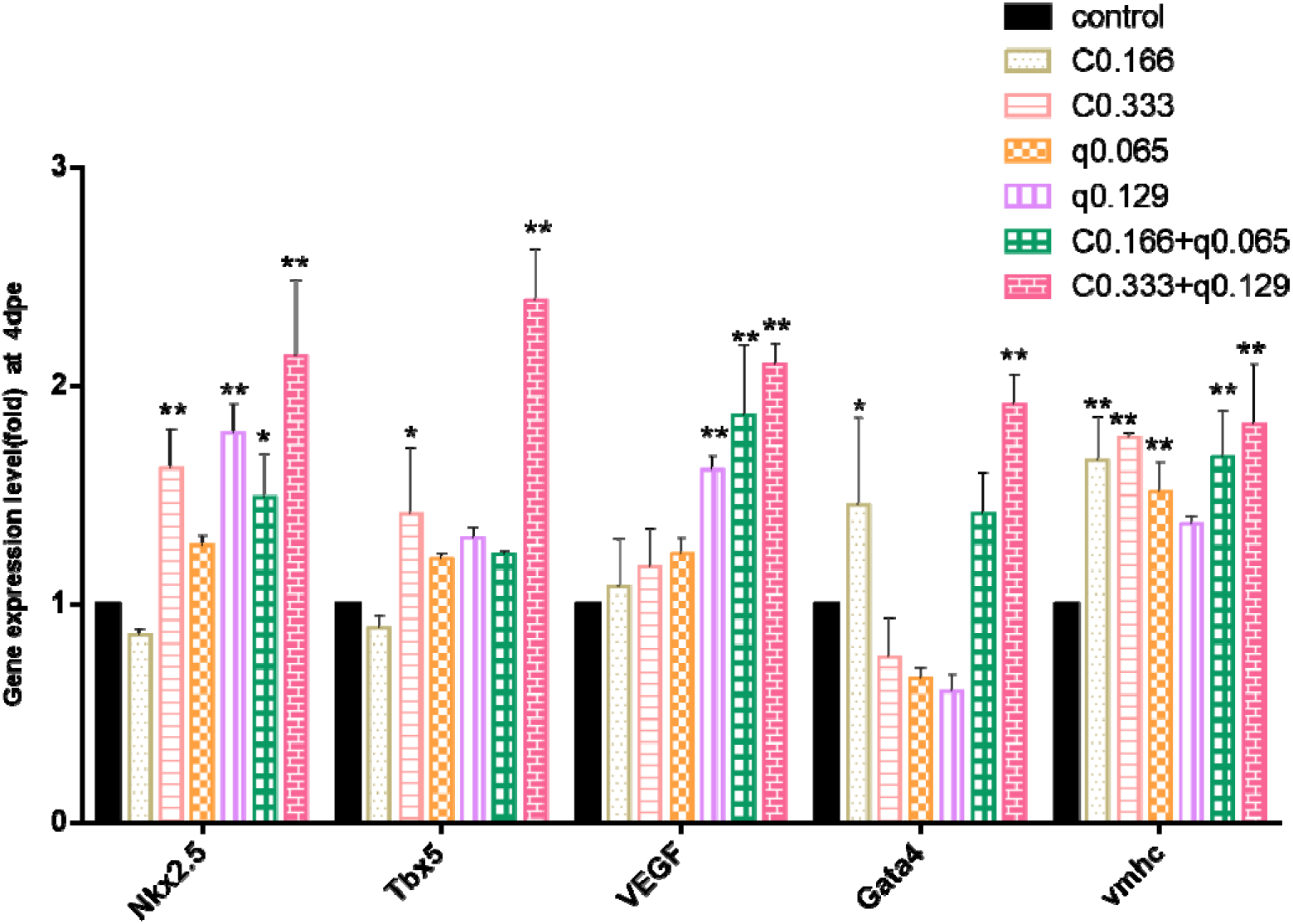
The expression level of key cardiac genes (*Nkx2.5*, *Tbx5*, *VEGF*, *Gata4*, *vmhc*) in control and treated zebrafish groups for 96 hpf. Results are presented as mean ± SD of three replicate samples (determined by Dunnett post-hoc comparison, ^◻^*p* < 0.05; ^◻◻^*p* < 0.01).

## 4. Discussion

In this study, the acute toxicity (96 hpf-LC_50_) of cyhalofop-butyl and quizalofop-p-ethyl to zebrafish were 0.637 and 0.248 mg/L respectively, which is consistent with previous result (Zhu et al., 2015; Zhu et al., 2022). In addition, the 96 h-LC_50_ of cyhalofop-butyl was 0.664 mg/L, and the 96 h-LC_50_ of quizalofop-p-ethyl was 0.259 mg/L under combined exposure. The LC_50_ values were 1.04 times of those under single exposure. The results calculated by the additive index method showed that the combined exposure (toxicity ratio 1:1) had an antagonistic effect on zebrafish embryos. However, different proportions of mixed pollutants will also produce different types of toxic effects.

Studies have shown that the combination of imidacloprid, acetochlor and tebuconazole has synergistic toxicity to zebrafish with the toxicity ratios of 1:2:2, 1:4:4, 2:4:1 and 4:1:4, while the combination of 1:1:1, 1:1:2, 2:1:2, 2:2:1 and 4:2:1 has antagonistic toxicity to zebrafish. Therefore, the toxic effects of cyhalofop-butyl and quizalofop-p-ethyl on zebrafish embryos under different toxic ratios need to be further studied. The mechanism of the effect of cyhalofop-butyl and quizalofop-p-ethyl on zebrafish embryos in the ratio of 1:1 is still unclear, so we explored the preliminary toxic mechanism of combined exposure on zebrafish embryos in the ratio of 1:1 antagonism. Cyhalofop-butyl can induce developmental toxicity, oxidative stress and apoptosis of zebrafish embryos (Zhu et al., 2015), while quizalofop-p-ethyl has developmental toxicity, cardiotoxicity and reproductive toxicity to zebrafish (Zhu et al., 2022; Zhu et al., 2017).

In this study, the results of joint developmental toxicity test showed that co-exposure caused a series of adverse effects during zebrafish embryo development, including abnormal voluntary movement, decreased heart rate, hatching inhibition, growth inhibition and various teratogenic effects, including yolk cyst and pericardial edema. Heart rate change is an important index to evaluate cardiac toxicity. Many studies have found that exposure to chemicals can affect the heart of zebrafish embryos. For example, study pointed that exposure to boscalid can cause the heart malformation of zebrafish embryos, and significantly inhibit the embryo heartbeat (Qian et al., 2018). It has been reported that difenoconazole can cause a lot of symptoms during the development of zebrafish embryos, such as slow heart rate, morphological abnormality and hatching inhibition (Mu et al., 2013). Dioxin and dioxin-like compounds can cause morphological abnormalities of zebrafish embryos, including pericardial edema and craniofacial abnormalities (Tokunaga et al., 2016).It was found that the heart function of zebrafish embryos was impaired, the heart rate decreased obviously, and serious changes such as pericardial edema, yolk sac edema and yolk sac deformation occurred during the time when zebrafish embryos were exposed to cyhalofop-butyl and quizalofop-p-ethyl, and the co-exposure aggravated these changes. In addition, abnormalities mainly occur in cardiac areas, such as pericardial edema. This finding indicated that abnormal yolk sac edema and pericardial edema may be caused by heart damage, and the combined exposure of cyhalofop-butyl and quizalofop-p-ethyl makes the heart damage more serious, which makes the effects of abnormal voluntary movement, decreased heart rate, hatching inhibition, growth inhibition and various teratogenic effects after combined exposure more severe.

Apoptosis is an active cell death process with obvious morphological characteristics and energy dependence (Abrahams et al., 2014). Many environmental pollutants, such as pesticides (Qian et al., 2018), heavy metals (Kp et al.), polycyclic aromatic hydrocarbons (Deng et al., 2009b), etc., disturb the expression of protein and nucleic acid in cells in the process of contacting organisms, thus inducing apoptosis. Therefore, taking apoptosis as a detection index can reflect the early toxic effect of pollutants on organisms, which is of great value for the evaluation of environmental pollutants. Apoptosis may lead to early developmental malformation. For example, zebrafish larvae exposed to hexabromocyclododecane (HBCD) were triggered by apoptosis genes, which led to the increase of deformity and the decrease of survival rate (Deng et al., 2009a). After being exposed to boscalid, zebrafish embryos suffered from developmental toxicity, such as slow heartbeat, cell edema and spinal deformation, through apoptosis and lipid metabolism (Qian et al., 2018). In this study, the single or combined exposure of cyhalofop-butyl and quizalofop-p-ethyl caused a series of gene expressions related to apoptosis in zebrafish embryos, such as *bax*, *bcl-2*, *p53*, *mdm2*, *caspase9*, etc., and the activities of caspase-3 and caspase-9, the important executors of apoptosis, were also significantly induced. This conclusion proves that apoptosis can induce a series of adverse effects (heart rate drop and deformity, etc.) during embryo development. The expression of apoptosis-related genes is related to the process of apoptosis signal pathway, such as cytochrome C, *bcl2*, *bax*, *p53*, *apaf1* (Zhao et al., 2009). Cytochrome C was released from mitochondria to cytoplasm during apoptosis stimulation (Hildeman et al., 2003). The activation of *p53* expression indicates that the cells are in an apoptotic state (Jin et al., 2012). In addition, Bcl-2 family proteins are composed of anti-apoptosis (e.g., bcl-2) and pro-apoptosis members (e.g., bax), which play an important role in inhibiting or promoting apoptosis mainly through mitochondrial pathway (Hildeman et al., 2003; Li et al., 2009). The activation of *caspase-9* is triggered by the release of cytochrome C in mitochondria and its interaction with *apaf1*(Pallardy et al., 1999; Yoshida et al.), and the activated *caspase-9* will further activate regulatory factors such as *caspase3* downstream (Lindenboim et al., 2000).

In this study, when cyhalofop-butyl (0.166 and 0.333 mg/L) and quizalofop-p-ethyl (0.065 and 0.129 mg/L) were exposed alone, we observed that *p53*, *bax*, *bcl-2* and *apaf1* were up-regulated with the increase of exposure concentration. Compared with the control group, the gene transcription of *caspase-3* and *caspase-9* was also up-regulated, and *caspase-9* was significantly up-regulated in quizalofop-p-ethyl treatment group. In the co-exposure (0.166+0.065 mg/L, 0.333+0.129 mg/L) treatment group, we observed that *p53*, *bax*, *bcl-2*, *apaf1*, *caspase-3* and *caspase-9* were up-regulated with the increase of exposure concentration, and up-regulated significantly in the highest concentration (0.333+0.129 mg/L). We also detected the apoptosis induced by zebrafish embryos in vivo by caspase activity assay. It was found that the gene expression and activity of caspase-3 and caspase-9 increased when exposed to cyhalofop-butyl and quizalofop-p-p-ethyl alone or in combination, and significantly increased in the co-exposed highest concentration (0.333+0.129 mg/L) treatment group. It is suggested that cyhalofop-butyl and quizalofop-p-p-ethyl may induce mitochondrial cytochrome C release by up-regulating pro-apoptotic factor *bax*, *caspase-3* and *caspase-9* are important executors of inducing apoptosis. On the other hand, it can induce the initiation of *p53* pathway and activate the apoptosis of zebrafish embryos. Subsequently, these enzymes promote cell apoptosis, resulting in pericardial cyst and yolk cyst, which in turn affects the heart function, and eventually leads to embryo death.

As the first organ in zebrafish embryo development (Bakkers, 2011), the formation of heart is a very complicated process (Gonzalez-Rosa et al., 2017). Many key genes, such as Tbx5 (Ingham, 2000), Nkx2.5 and gata4 (Välimäki et al., 2017), play an important role in the development and maturation of zebrafish heart. In this study, the exposure of two drugs will cause obvious cardiotoxicity to zebrafish embryos. By observing the heart rate of zebrafish embryos, it was found that the heart beats of the embryos in the control group was regular, while the heart beat rhythm of the embryos in the treatment group was irregular, and the heart rate decreased obviously. And we found that cyhalofop-butyl and quizalofop-p-p-ethyl and binary mixed exposure could induce the expression of genes related to heart development of zebrafish embryos, such as *Nkx2.5*, *Tbx5* and *VEGF*. Compared with the control group, the expressions of *Tbx5* and *Gata4* genes were up-regulated under combined exposure, and the expressions of *Nkx2.5*, *VEGF* and *vmhc* were significantly up-regulated in the low concentration group (0.166+0.065 mg/L). Compared with the control group, the expression of these genes in high concentration group (0.333+0.129 mg/L) was significantly up-regulated. Compared with the single dose of cyhalofop-butyl and quizalofop-p-p-ethyl on the expression of heart development-related genes in zebrafish embryos, the co-exposure showed synergistic effect on the expression of *Nkx2.5*, *Tbx5*, *VEGF* and *vmhc*, and showed antagonistic effect on the expression of *Gata4*, which was significantly up-regulated with the increase of concentration. In addition, previous studies also found that cyhalofop-butyl exposure can produce zebrafish embryos with developmental abnormalities, which mainly include pericardial edema, yolk sac edema and abnormal yolk sac morphology, among which the most significant ones are pericardial edema and yolk sac edema (Zhu et al., 2015). After quizalofop-p-ethyl exposure, the embryo appeared pericardial edema, abnormal cardiac cyclization and atrial hypertrophy, and the expression of a series of genes (such as *gata4*, *Nkx2.5*, *Tbx5*, *VEGF*.) and proteins (Tbx5) related to heart development were induced to change (Zhu et al., 2022). This is consistent with our results. Generally speaking, quizalofop-p-ethyl, cyhalofop-butyl and co-exposure may induce the expression of genes related to heart development of zebrafish embryos, and further cause the malformation of zebrafish embryos such as hatching inhibition, heart rate decrease, autonomic motor inhibition and pericardial cyst. Therefore, when assessing the harm of pesticides to aquatic organisms, the combined effect of combined pollution should be fully considered.

## 5. Conclusions

In summary, our results showed that the single or combined exposure of cyhalofop-butyl and quizalofop-p-p-ethyl in water had significant negative effects on the development of zebrafish embryos. During the development of zebrafish embryos, obvious developmental effects such as hatching inhibition, decreased heart rate, shortened body length and deformity were observed. The effects of apoptosis and heart development may be the responsibility of the toxicity of combined exposure to zebrafish embryo development. Aquatic organisms are often exposed to a variety of compounds, and a variety of pesticides combine to produce different toxic effects. Therefore, it is of great significance to study the combined effects of combined pollution when assessing the harm of pesticides to aquatic organisms.

## Acknowledgments

This work was supported by the Basic and Advanced Research Project of Chongqing (cstc2022jcyjAX0532).

## Conflict of interest

The authors declare that they have no known competing financial interests or personal relationships that could have appeared to influence the work reported in this paper.

## Notes

### Competing Interest Statement

The authors have declared no competing interest.

