## Supplementary Information for "Combined Developmental Toxicity Of Cyhalofop-butyl And Quizalofop-p-ethyl On The Zebrafish (Danio rerio) Embryos"

### Summary of Supplymentary Information

**Table:**

Table S1. Sequences of primer pairs used in the real-time quantitative PCR reactions.

**Table S1** Sequences of primer pairs used in the real-time quantitative PCR reactions.

| Target gene | Primer sequences | Accession number |
|---|---|---|
| *p53* | Forward-GGGCAATCAGCGAGCAAA | AF365873.1 |
| | Reverse-ACTGACCTTCCTGAGTCTCCA | |
| *bax* | Forward-GGCTATTTCAACCAGGGTTCC | AF231015.1 |
| | Reverse-TGCGAATCACCAATGCTGT | |
| *bcl-2* | Forward-AGGAAAATGGAGGTTGGGATG | NM_001030253.2 |
| | Reverse-TGTTAGGTATGAAAACGGGTGGA | |
| *apaf-1* | Forward-TTCTACAGTAAACGCCCACC | AF251502.1 |
| | Reverse-TATCTAGTATTTCCCCATATTCC | |
| *Caspase-9* | Forward-AAATACATAGCAAGGCAACC | NM_001007404.2 |
| | Reverse-CACAGGGAATCAAGAAAGG | |
| *Caspase-3* | Forward-CCGCTGCCCATCACTA | NM_131877.3 |
| | Reverse-ATCCTTTCACGACCATCT | |
| *β-Actin* | Forward-CGAGCAGGAGATGGGAACC | AF057040.1 |
| | Reverse-CAACGGAAACGCTCATTGC | |
| *GATA4* | Forward- CCAGACACACACCAGCTCTACAC | NM_131236.1 |
| | Reverse- ATCAGGCTGTTCCACACTTCACT | |
| *NKX2.5* | Forward- TAGTAATTTTCCGAGTCCAGGC | NM_131421.1 |
| | Reverse- GCACGTTATTTAGATCCCCAAC | |
| *TBX5* | Forward- CGCTATAAATTCGCCGATAACAA | NM_130915.1 |
| | Reverse- AGACACCAGTTGCCTCATCCAG | |
| *VEGF* | Forward-TGCCCACATACCCAAAGAAGG | NM_131408.3 |
| | Reverse-CACAGCGCATGAGAACCACA | |
| *vmhc* | Forward-ACATAGCCCGTCTTCAGGATTTGG | NM_001077464.2 |
| | Reverse-GAGAGAAAGGCAAGCAAGTACTGG | |
